# Pharmacogenetic variation in Neanderthals and Denisovans and implications for human health and response to medications

**DOI:** 10.1101/2021.11.27.470071

**Authors:** Tadeusz H. Wroblewski, Kelsey E. Witt, Seung-been Lee, Ripan S. Malhi, Emilia Huerta-Sanchez, Fernando Villanea, Katrina G. Claw

## Abstract

Modern humans carry both Neanderthal and Denisovan (archaic) genome elements that are part of the human gene pool and affect the life and health of living individuals. The impact of archaic DNA may be particularly evident in pharmacogenes – genes responsible for the processing of exogenous substances such as food, pollutants, and medications. However, the health implications and contribution of archaic ancestry in pharmacogenes of modern humans remains understudied. Here, we explore eleven key cytochrome P450 genes (*CYP450*) involved in drug metabolizing reactions in three Neanderthal and one Denisovan individuals as well as examine archaic introgression in modern human populations. We infer the metabolizing efficiency of these eleven *CYP450* genes in archaic individuals and find important phenotypic differences relative to modern human variants. We identify several single nucleotide variants shared between archaic and modern humans in each gene, including some potentially function-altering mutations in archaic *CYP450* genes, which may result in altered metabolism in living people carrying these variants. We highlight three genes which show evidence for archaic introgression into modern humans, as well as one additional gene that shows evidence for a gene duplication found only in Neanderthals and modern Africans.

## Introduction

The cytochrome P450 (*CYP450*) genes encode oxidase enzymes that function in metabolism of endogenous small molecules and in detoxification of exogenous (or xenobiotic) compounds. This gene family is present in all mammals, with 57 active genes and 58 pseudogenes coding for CYP450 enzymes in humans ^1^. These xenobiotic-substrate enzymes have been studied extensively because of their roles in the absorption, distribution, metabolism, and excretion (ADME) of pharmaceuticals and drug development. The evolution of xenobiotic CYP450 enzymes may be driven by organisms’ need to metabolically detoxify foreign compounds - often toxic chemicals produced by plants, fungi, and bacteria - in the local environment. The *CYP450* genes show evidence of positive selection and high allele frequency variation in humans ^1, 2^. It has been suggested that the shift from hunting and gathering to food production in humans may have profoundly changed the selective effect of certain CYP450 enzymes ^2, 3^. Although these genes have been studied extensively in humans and other mammals, the extent of genetic variation in archaic individuals and its relation to modern human variation has not been addressed. Genetic variation in ADME genes varies extensively across modern human populations, and identifying alleles inherited through archaic introgression may be informative about the origin of specific pharmacogenetic (i.e., genes that modulate the body’s response to pharmaceuticals; PGx) variants and their phenotypes, including metabolizer status.

Throughout human evolutionary history, our species has adapted to numerous distinct and varied challenging environments, which present the need to metabolize new xenobiotic substances, such as food, pollutants, and, most recently, medications ^4^. Modern humans encountered new environmental challenges as they dispersed first throughout and then outside of the African continent, but they also encountered other hominin species already adapted to those novel environments (i.e., Neanderthals and Denisovans) ^5, 6^. The direct sequencing of multiple Neanderthal and Denisovan genomes has revealed a complex history of admixture between these archaic humans and the ancestors of modern humans ^7, 8^. Most modern humans carry a small but significant portion of archaic ancestry, which has been targeted by natural selection ^9^. Purifying selection — or negative natural selection — has removed many archaic variants in functional genomic regions ^10–12^, while some archaic variants may have been lost through genetic bottlenecks or drift in modern human populations. In spite of selective pressures, there are still functional regions for which archaic variants are found at very high frequency in living humans through the effects of positive natural selection ^13–16^. One of the most well-known examples of adaptation through archaic admixture is high-altitude adaptation in Tibetans, a population in which the Denisovan variant of the gene *EPAS1* is found at extremely high frequency, and has conferred them with resistance to hypoxic stress in high-altitude environments ^14^.

The *CYP450* genes offer a promising target for adaptation through archaic introgression as humans encountered novel exogenous substances as they expanded their range. Neanderthals and Denisovans may have possessed *CYP450* variants fine-tuned to metabolizing substances found in their native habitats of Eurasia and Siberia. As modern humans moved outside of Africa, they likely faced novel environmental factors which may have influenced selective pressures; therefore, advantageous archaic *CYP450* variants introduced to modern humans through admixture may have been retained in the modern human gene pool by natural selection.

Here, we examine genetic variation and predict metabolizer phenotypes in eleven *CYP450* genes in three Neanderthal and one Denisovan individuals using publicly available high-resolution whole genome sequencing data. These genes represent enzymes which are responsible for up to 75% of the metabolism of commonly prescribed drugs and other xenobiotic substrates ^17–20^, which could have implications to modern human health, disease, and drug safety and efficacy. In addition, we examine the patterns of allele sharing between archaic and modern humans to identify haplotypes that may have been introgressed. This study aims to increase our understanding of introgressed genetic variants and the different selective regimes and environments acting on these enzymes.

## Subjects and Methods

### Samples

We investigate population-specific *CYP450* gene variation by combining data from the 1000 Genomes Project ^21^, the Simons Genome Diversity Project ^22^, the complete Neanderthal genomes ^23–25^, and the Denisovan genome ^26^. We extracted coding variation from four archaic human genomes: the Denisovan individual from the Denisova cave in the Altai Mountains (∼21X coverage), a Neanderthal individual from Croatia (∼42X coverage), and two Neanderthal individuals from the Altai Mountains: one from the Denisova cave (∼30X coverage), and one from the Chagyrskaya cave (∼28x coverage). These individuals are estimated to be at least 50,000 years old. We additionally utilized a set of seventy published chimpanzee genomes ^27^, which were used as a proxy for variation that is ancestral to modern and archaic humans.

### Variant Calling

Sequencing data for the three Neanderthal individuals and one Denisovan individual are publicly available with alignment and processing of the BAM files previously outlined ^23–26^. The BAM files were used to calculate depth of coverage in the sample’s genomic regions (see section *Star Allele Calling*), and then processed for variant calling. *De novo* variant calling was performed from BAM files instead of using the previously generated variant call files, which were likely more stringent and may have removed heterozygous sites. Sample BAM files were separated by chromosome for easier downstream processing using samtools (version 1.10 with hts lib 1.10) ‘view’ function ^28^. Variant calling was performed for each *CYP450* gene of interest, using PyPGx (version 0.1.37) ‘bam2vcf’ which implements the Genome Analysis Toolkit (GATK, version 4.1.9.0) ‘HaplotypeCaller’ function (options -emit-ref-confidence GVCF; - minimum-mapping-quality 10) followed by GATK ‘GenotypeGVCF’ to merge individual samples ^29^. Variants were called against the HumanG1Kv37 hg19 reference assembly. GATK ‘VariantFiltration’ was additionally utilized to annotate variants with a quality score of 50. Variants from modern human individuals from the 1000 Genomes Project ^21^ included for further analysis were called using the same procedure.

For individual SNV analysis, variant call format (VCF) files were further filtered by the phred-scaled quality score (QUAL) > 40 and gene regions were determined as the RefSeq start and end coordinates of the gene with 2000 base pairs upstream to account for the promoter region using bcftools (version 1.10.2 with hts lib 1.10.2) ^28, 30^. Variant read depth was calculated with the PyPGx (version 0.14.0, https://pypgx.readthedocs.io/en/latest/readme.html) pipeline (described below). Variants were additionally filtered for polymorphic sites that had a BAM read depth ≤ 1 for one allele or a ratio of minor allelic read depth to major allelic read depth < 0.2.

Variant annotation was conducted through ANNOVAR (-protocol refGeneWithVer, knownGene, avsnp150, ljb26_pp2hvar) which utilizes dbSNP150, RefSeq, and the UCSC genome browser to identify and characterize known variants; the occurrence of novel SNVs were additionally confirmed with Gnomad (v2.1.1) ^30–34^. Variant locations were identified with ANNOVAR, which annotates gene location based on RefSeq and UCSC genome browser data. Variants are defined as exonic, splicing, ncRNA, 3’ and 5’UTR, intronic, or in the promoter region. If the variant type differed between the RefSeq and UCSC gene annotation, the called location is determined by an ordering precedence: exonic/splicing > ncRNA > 3’/5’UTR > intronic > promoter region.

Potentially function-altering variants were identified using three algorithms to rank variant deleteriousness: Combined Annotation Dependent Depletion (CADD) integrates multiple weighted metrics to identify deleterious variants ^35, 36^, Sorting Intolerant From Tolerant (SIFT) predicts the functional impact of amino acid substitution caused by genetic variation ^37^, and Polymorphism Phenotyping (PolyPhen) identifies the predicted structural and functional outcome of amino acid substitution ^38^. Variants considered potentially function-altering (referred to as “damaging” or “pathogenic” by algorithms) were defined as SNVs that are in the top 1% of deleterious variants by CADD score (Phred-normalized CADD score ≥ 20), deleterious by SIFT prediction (SIFT score ≤ 0.05) or damaging by Polyphen2 prediction (Polyphen2 score between 0.15-1.0 as possibly damaging and 0.85-1.0 as probably damaging).

### Star Allele Calling

*CYP450* “star alleles” (phased haplotypes) were called using Stargazer ^39, 40^ and PyPGx (version 0.14.0), bioinformatics tools for identifying star alleles by detecting single nucleotide variants (SNVs), insertion/deletions (indels), and structural variants (SVs) of pharmacogenes. The star allele system is a nomenclature used specifically for pharmacogenes ^41^ – star alleles refer to haplotype patterns in pharmacogenes and are usually associated with protein activity levels. Phased haplotypes were called for eleven pharmacogenes that represent enzymes which are responsible for up to 75% of the metabolism of commonly prescribed drugs ^17^: *CYP1A2*, *CYP2A6*, *CYP2B6*, *CYP2C19*, *CYP2C8*, *CYP2C9*, *CYP2D6*, *CYP2E1*, *CYP2J2*, *CYP3A4*, and *CYP3A5.* VCF files, without additional QC filtering, along with the read depth data for each gene were the input into the haplotype callers, which utilize the statistical phasing software BEAGLE to phase the variants into haplotypes ^42^. Star alleles were then inferred by identifying the haplotype calls from core variants of each star allele, as well as tag variants, which define sub-alleles of each star allele. Star allele diplotype calls were considered “non-determinant” if any variant used in star allele detection method had a read depth ≤ 1.

SV calls were assessed with PyPGx (version 0.14.0) using three control genes: vitamin D receptor (*VDR)*, epidermal growth factor receptor (*EGFR)*, and ryanodine receptor 1 (*RYR1)*. Depth of coverage data was calculated using PyPGx to call the read depth at each position and was compared to the control genes, which are stable read regions that are highly conserved and do not fluctuate in copy number, to assess copy number variation in the eleven pharmacogenes^43^. Final calls for star alleles were determined as the call that was consistent in the highest number of control genes. Phenotype prediction was conducted in PyPGx by converting the called diplotype into an activity score, which is then used to predict the gene phenotype ^39, 41^. Calls were additionally confirmed with visual inspection of read depth regions for the three control genes.

### Identifying archaic alleles in modern populations

To identify the presence of archaic *CYP450* variants in modern human populations, we calculated the allele frequency of the archaic variants described above in modern humans. We used modern human genomes from all populations sequenced for the 1000 Genomes Project ^21^, as well as the Papuans sequenced as part of the Simons Genome Diversity Project ^22^. We also focused on a subset of SNVs that were more likely to be introgressed, which are found outside of Africa but not in African populations. These SNVs had to have a frequency in African populations of less than 1 percent and be present in at least one non-African population with an allele frequency greater than 1 percent. For all results described in the text, we used the archaic VCF generated using the methods in this paper for comparison with modern humans. For comparison, we repeated these analyses with the published VCFs that are associated with the original archaic genome sequencing studies ^25, 26^. Alleles were categorized as “derived” or “ancestral” based on the ancestral allele calls identified by the 1000 Genomes Project ^18^. We additionally examined these pharmacogenes in a cohort of seventy publicly available chimpanzee genomes ^27^ to determine if any of the variants shared between archaic and modern humans might be found in chimpanzees as well, representing shared ancestral variation.

We used Haplostrips ^44^ to calculate the distance between haplotypes of all *CYP450* genes in modern humans in the 1000 Genomes Panel relative to a reference haplotype. Distances in Haplostrips are Manhattan distances - simply the number of SNVs with different alleles between the two sequences. Haplotypes are re-ordered by decreasing similarity with the archaic reference haplotype. In this case, the Haplostrip is polarized to a Vindija Neanderthal reference haplotype (a consensus of the two archaic chromosomes) or a Denisovan reference haplotype, and each subsequent haplotype is ordered by genetic similarity, from most related to least related. For this analysis, we looked at haplotypes of each *CYP450* gene at the gene coordinates, plus 5000 base pairs downstream and upstream to capture more linked neutral variation and compared the 1000 Genomes individual’s haplotypes with archaic haplotypes.

For one pharmacogene, *CYP2B6*, we observed highly divergent haplotypes found in the Vindija Neanderthal and a small number of African individuals (*n =* 11). To determine how the divergent Vindija *CYP2B6* haplotype compared to the divergence between the Vindija Neanderthal and other Neanderthals in other regions of the genome, we calculated the pairwise distance between the Chagyrskaya Neanderthal and Vindija Neanderthal genomes with 29.1 kb windows (*CYP2B6* gene length) across the genome. To explore SNV sharing between the *CYP2B6* haplotypes, we divided the archaic and modern humans into four groups: the Vindija genome (containing the divergent haplotype), the other three archaic genomes, the eleven Africans with the divergent haplotypes, and the rest of the modern Africans without the divergent haplotypes. Each group was scored for the presence or absence of a given SNV, and the sharing of these SNVs was summarized in a Venn Diagram (Figure S3).

### ILS Calculation

While shared ancestry between archaic humans and modern humans outside of Africa may suggest that introgression has occurred, this pattern may also represent Incomplete Lineage Sorting (ILS), in which a haplotype that existed in an ancestral population (for example, the shared ancestor of humans and Neanderthals) is lost from one or more descendant populations (for example, in Africans). To attempt to distinguish between the two possibilities, we assessed the probability of ILS occurring in a given region using a previously published equation ^14^:

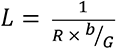

Briefly, the expected length of a shared ancestral sequence haplotype (L) is calculated as the inverse of regional recombination rate (R) multiplied by the branch length (b) over the generation time (G) The probability (p) of seeing a shared haplotype of length (m) follows a Gamma distribution with a shape (S) of 2 and a rate (r) of 1/L:

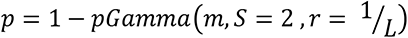

Here, we determined the regional recombination rates (R) using HapMap ^45^, used a generation time (G) of 29 years ^46, 47^, and assumed a branch length (b) of 550,000 years with a 50,000 year timeframe of Neanderthal-human interbreeding ^23, 47^.

### Data Visualization

All data visualization was conducted in R Studio Suite (R version 4.0.2, https://www.r-project.org/) with the use of the CRAN library *ggplot2* for graphical visualization of variant frequency, star allele diplotype calls, deleterious variant occurrence, and haplotype distance visualization ^48, 49^. The R package *venneuler* was used to create the Venn diagram in Supplemental Figure 3 (https://CRAN.R-project.org/package=venneuler).

## Results

### CYP450 Variation in Neanderthals and Denisovans

In this study, we investigated genetic variation in eleven *CYP450* genes: *CYP1A2, CYP2A6, CYP2B6, CYP2C8, CYP2C9, CYP2C19, CYP2D6, CYP2E1, CYP2J2, CYP3A4,* and *CYP3A5*. We identified a total of 1,623 single nucleotide variants (SNVs) across the eleven *CYP450* genes investigated in one Denisovan (from Denisova Cave), and three Neanderthals (from the Vindija, Denisova Cave/Altai, and Chagyrskaya sites). Of the variants identified in the four archaic individuals, the majority were intronic or non-coding, while 6.8% (*n=*111) were in promoter regions and 4.7% (*n=*77) were exonic (Figure 1A). A summary table of genetic variants across gene and location is shown in Table S1.

**Figure 1.**
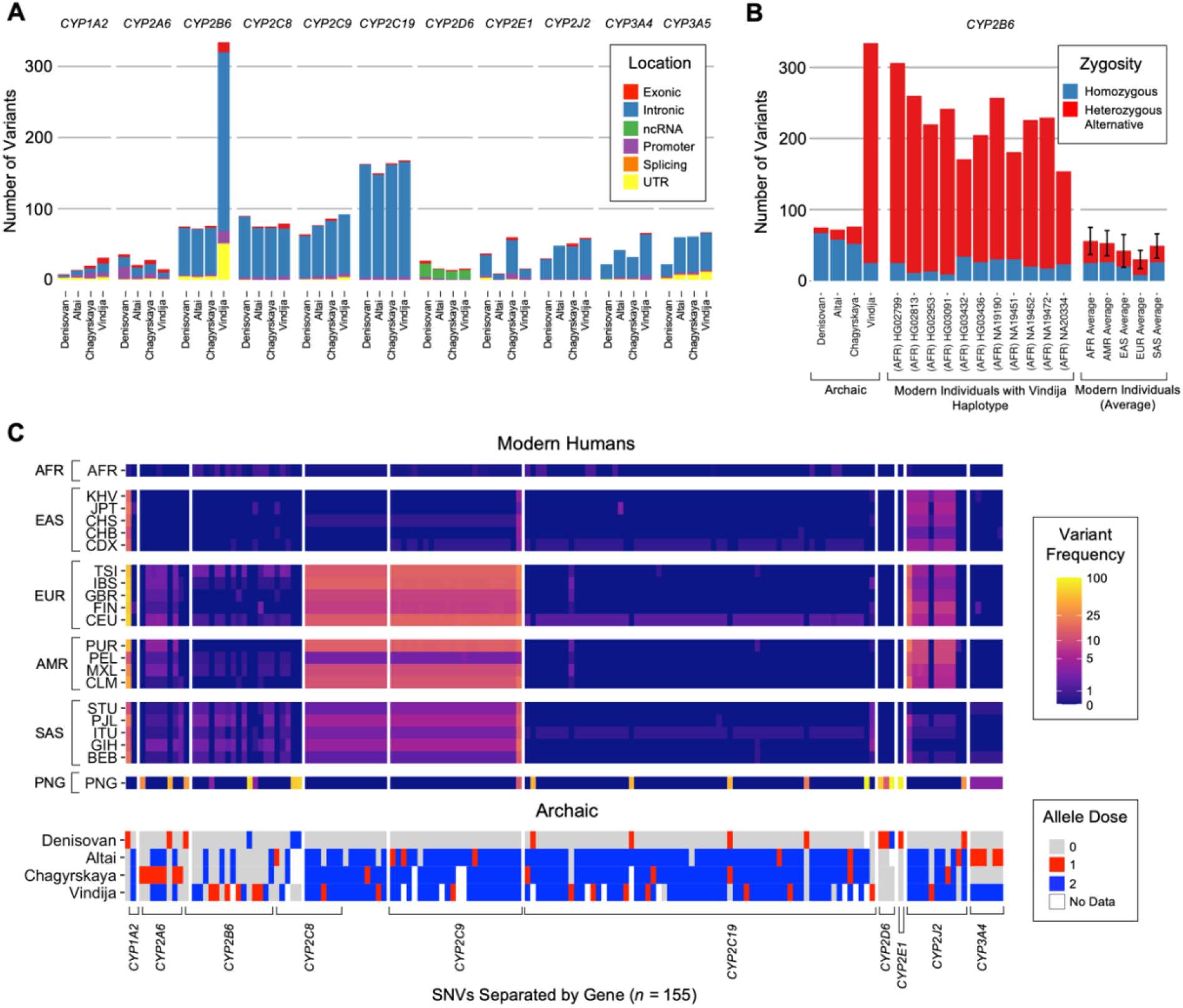
*CYP450* genetic variation in archaic individuals and shared with modern humans. (A) The total number of SNVs identified in each archaic individual displayed as a stacked bar plot. Each color displays the relative variant location identified using ANNOVAR with 3’ & 5’ untranslated regions grouped together (UTR), and the promoter region (Promoter) characterized as any upstream variants that are within 2000 bp of the gene. (B) A larger number of variants were identified in *CYP2B6* in the Vindija Neanderthal. Bar plots have been faceted by the archaic individuals, the modern individuals identified to share the Vindija *CYP2B6* haplotype, and the average number of variants for 10 modern individuals from each super population of the 1000 Genomes Project. Variants are colored by zygosity (i.e., heterozygous if the variant site has one alternative allele and homozygous alternative if the variant site has two alternative alleles at that position). The height of each stacked bar plot represents the total number of variants for each individual and the number of variants for the modern individuals have been reported as the mean with error bars representing the standard deviation. (C) A subset of archaic variants (*n =* 155) was identified to have an allele frequency <1% in African populations and >1% in at least one non-African population. Sub-populations have been grouped as African (AFR), East Asian (EAS), European (EUR), admixed American (AMR), South Asian (SAS), and Papuan (PNG) and allele frequencies for each population are shown on the gradient heatmap. The allele dose of each archaic variant is shown in the heatmap on the bottom, with an allele dose of 0 representing homozygous reference and an allele dose of 2 representing homozygous alternative.

The Vindija Neanderthal harbored a particularly large number of SNVs in the *CYP2B6* gene, which was an outlier relative to other Neanderthals and warranted further scrutiny. The Vindija Neanderthal had 334 variant sites in *CYP2B6* with approximately 92% of these variant sites being heterozygous (Figure 1B), while the other three archaic genomes had fewer than 100 variants each (Denisovan: *n=*75; Altai: *n=*72; Chagyrskaya: *n=*76). Given that *CYP2B6* shares significant homology with the pseudogene *CYP2B7,* which is located nearby (40.6 kb) ^50^, we assessed the read depth at the *CYP2B6* locus in the Vindija Neanderthal genome to ensure that the elevated variation identified was not a result of read mis-mapping with the paralog gene (Figure S1A). The *CYP2B6* locus in the Vindija Neanderthal was also identified as highly divergent from other Neanderthals relative to the rest of the genome and is within the top 1% of ∼30-kb windows based on divergence between the Chagyrskaya and Vindija Neanderthals (pairwise distance = 0.00175, Figure S2).

To further assess the genomic landscape of the archaic individuals in the context of modern humans, we compared the archaic *CYP450* genetic variation to the variation in modern human individuals from the 1000 Genomes Project ^21^. Interestingly, a subset of modern individuals carried a haplotype related to the Vindija Neanderthal *CYP2B6* gene and present a similar increase in variation to the Vindija Neanderthal (Figure 1B). The eleven modern human individuals identified that carry this haplotype were all from African populations (ASW, ESN, GWD, LWK, MSL, YRI), and the haplotype is absent outside of Africa. These individuals were additionally assessed for read mis-mapping (Figure S1B), and demographic information is described in Table S2. The African divergent haplotype and the Vindija Neanderthal divergent haplotype uniquely share some SNVs, but they also each have a large number of variants that are exclusive to their individual haplotype (Figure S3).

Across the eleven *CYP450* genes investigated, archaic variation present in modern humans from the 1000 Genomes Project was identified as SNVs that were shared between modern and archaic humans, while found at a frequency of less than 1% in African populations, which we refer to as ‘archaic SNVs’ hereon. We identified a total of 155 archaic SNVs that fit these criteria, including 41 found in human populations at a frequency of 10% or higher, and four that were exonic (Figure 1C, full list of variants in Table S3). The majority of these 155 SNVs are derived (91.0%) and only 7 of them (4.5%) were identified in the sample of chimpanzees we used for comparison, five of which were homozygous in all chimpanzees sampled (Table S3).While modern Papuans had fewer alleles shared with archaic humans, those that were shared were present at a much higher frequency and many of the variants were unique to Papuans, including all but one of the Denisovan-unique variants (Figure 1C, Table S3).

### Phased Diplotypes and Predicted Metabolizer Phenotypes

We identified the star allele composition of the eleven *CYP450* genes in archaic individuals using variant and depth of coverage data with previously validated methods ^39, 40^. The phased diplotype for each archaic individual was identified as the two primary haplotype calls based on the presence or absence of star allele-defining nucleotide and structural variants (Figure 2, Table S4). It is important to highlight that the haplotype callers utilized only assess nucleotide and structural variants included in previously identified star alleles, and therefore excludes novel pharmacogene haplotypes that may be present in these archaic humans.

**Figure 2.**
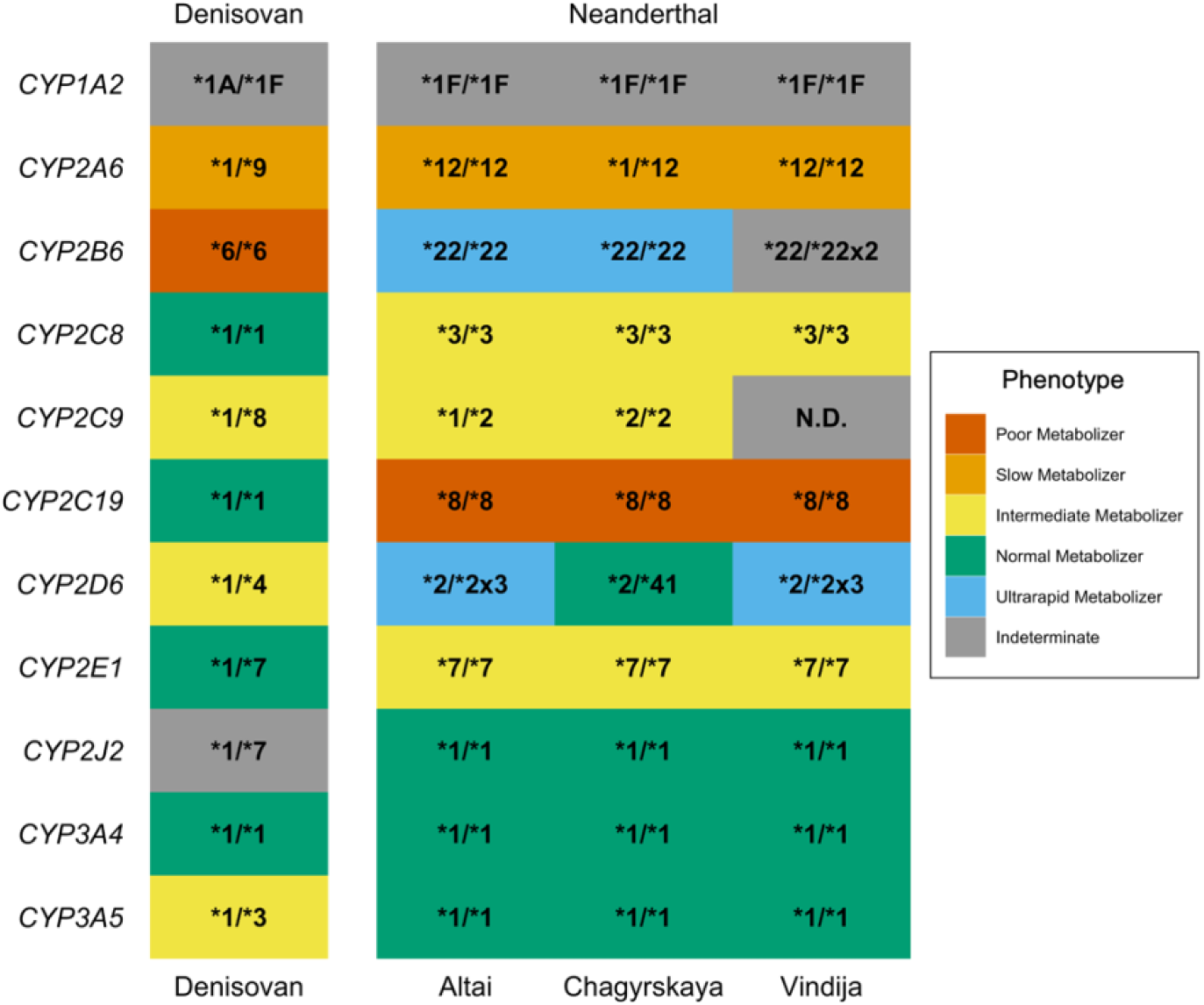
Phased diplotypes and predicted metabolizer phenotypes identified in archaic individuals. Heatmap showing the primary star allele diplotypes for each gene from the three Neanderthal and one Denisovan individuals. Each diplotype was determined as the two phased haplotypes, which are detected with the variant and read data from each gene. The fill of each tile represents the predicted phenotype determined by the combined activity score associated with the presented diplotype as identified in modern humans. Unknown phenotypes are represented with grey tiles and indicate that the functional effect of the diplotype has not been determined. A diplotype was considered non-determinant (N.D.) if there was a lack of read coverage at specific variant locations.

Pharmacogene diplotypes were identified for all eleven of the pharmacogenes studied for all four archaic individuals, with the exception of the Vindija diplotype for *CYP2C9*, which was indeterminate due to very low read coverage at informative variant sites (Figure 2). For each diplotype, the predicted metabolizer status phenotype was assessed as designated by the Pharmacogene Variation Consortium (https://www.pharmvar.org/) ^41^. We observed variability in the metabolic rate of the archaic pharmacogenes, ranging from “ultrarapid” (i.e., the enzyme would break down compounds much faster than expected) to “poor” (i.e., the enzyme is virtually non-functional) metabolizers (Figure 2). One nuance of this analysis is that the impact of being a slower or faster metabolizer is not necessarily universally positive or negative but is context-dependent and depends on multiple factors, including whether breaking down a compound activates or inactivates its function, as well as environmental influences and the genetic background of the individual bearing the diplotype. Additionally, the metabolizer phenotypes were established in modern humans, and so the phenotypes may slightly differ for the archaic individuals.

Structural variation (SV) was identified in three of the eleven pharmacogenes investigated– *CYP2A6, CYP2D6,* and *CYP2B6* (Figure 3). The Altai and Vindija Neanderthals presented the homozygous partial deletion hybrid *CYP2A6*12* variant (*\*12/*12*, Figure 3A), while the Chagyrskaya Neanderthal was heterozygous for the *\*12* haplotype (*\*1/*12*, Figure 3B). No structural variation was identified for the Denisovan individual in *CYP2A6* (Figure 3C). For *CYP2D6*, we identified a gene duplication event in the Altai and Vindija Neanderthals, presenting a *\*2/*2x3* diplotype (Figure 3D), while no structural variation was identified in the Chagyrskaya Neanderthal and Denisovan individuals (Figure 3E).

**Figure 3.**
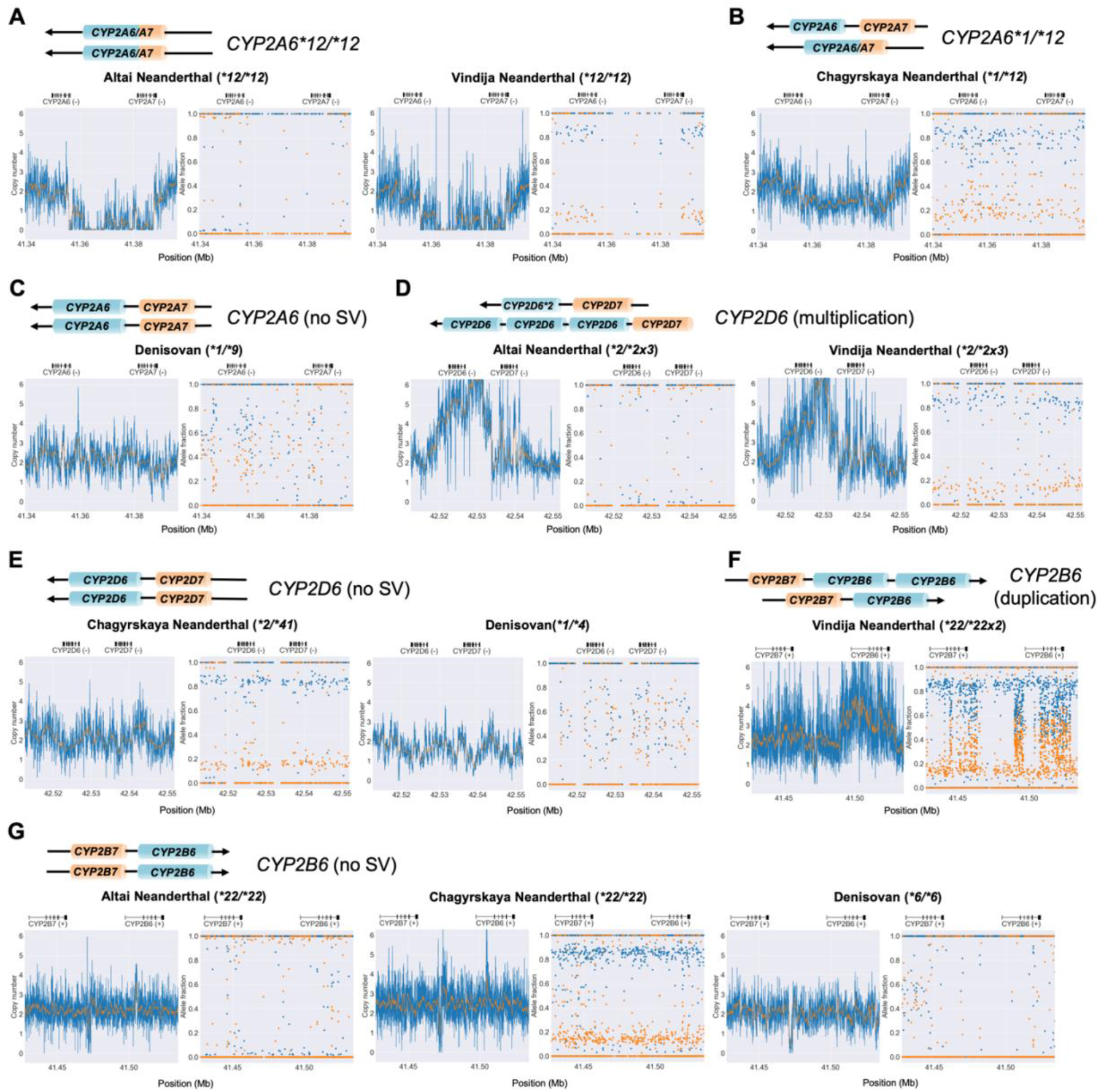
Structural variants detected in *CYP2A6*, *CYP2D6,* and *CYP2B6*. Copy number variation plots (left) demonstrating the structural variation and allele fraction plots (right) showing allelic fraction for each Neanderthal and Denisovan individual across *CYP2A6, CYP2D6,* and *CYP2B6*. The *CYP2A6/A7* structural variant, *\*12*, identified for the (A) Altai Neanderthal and Vindija Neanderthal which both present a *\*12/*12* diplotype, characterized by a partial deletion hybridization event in both gene copies. (B) The Chagyrskaya Neanderthal demonstrated a *\*1/*12* diplotype and (C) the Denisovan individual presented no structural variation. In *CYP2D6*, (D) the Altai and Vindija Neanderthals presented a gene multiplication while the (E) Chagyrskaya Neanderthal and Denisovan individual presented no structural variation in *CYP2D6*. For *CYP2B6*, (F) the Vindija Neanderthal displayed a gene duplication event, while (G) no structural variation was identified for the Altai Neanderthal, Chagyrskaya Neanderthal, and Denisovan individual. Copy number plots are estimated based on read data (displayed as the copy number normalized to the *VDR* control gene region) with the orange line indicating the copy number assessment for each gene. Allele fraction plots display the allele frequency of each variant for the two haplotypes, which demonstrates the allelic decomposition after identifying structural variation. For each structural variant, there is a schematic representation of the predicted structure of each gene, with an arrow indicating the direction of transcription. The hg19 genetic coordinates are presented on the *x axis* with the gene regions indicated directly above.

Interestingly, a gene duplication event in *CYP2B6* was identified in the Vindija Neanderthal (*\*22/*22x2*, Figure 3F-G) which was previously recognized as having a large number of heterozygous variants, as compared to the other Neanderthals and Denisovan for this gene (Figure 1B). Inspection of read depth in this region (Figure S1A) and allelic imbalance of *CYP2B6* for the Vindija Neanderthal (i.e., an allele fraction of 0.33 and 0.66, Figure 3F) further substantiates a gene duplication event of *CYP2B6* in the Vindija Neanderthal, as opposed to mis-mapping of reads containing *CYP2B7*. Perhaps most surprising is the fact that the group of eleven modern Africans with the similarly divergent haplotype (Figure 1B) also seem to share this gene duplication. Gene duplication was identified in eight of the 11 modern African individuals using PyPGx, and manual visual inspection suggests that the three other individuals also have elevated copy number in this region (Figure S4A). The duplicated *CYP2B6* gene is absent in the other archaic humans and in any non-African population in our dataset (Table S2).

### Potentially Function-altering SNVs

While haplotype calling methods can predict genotypes predicated on known star alleles, novel or un-classified variants are excluded from this method of analysis. Potentially function-altering SNVs were characterized using three programs that can predict functional impact of genetic variation: Combined Annotation Dependent Depletion (CADD v1.6), Sorting Intolerant From Tolerant (SIFT 4G v2.0.0), and Polymorphism Phenotyping (PolyPhen v2) ^35, 37, 38^. While “potentially damaging” is the standard terminology for SNVs that alter enzyme function, in this instance, these SNVs will be referred to as “potentially function-altering” as the impact of altered pharmacogenetic enzymes is dependent both on the substrate and on the individual’s genetic background. We identified 23 potentially function-altering variants in the eleven *CYP450* genes investigated (Figure 4), including 7 potentially novel variants that were not observed in any variant annotation databases utilized (see Methods).

**Figure 4.**
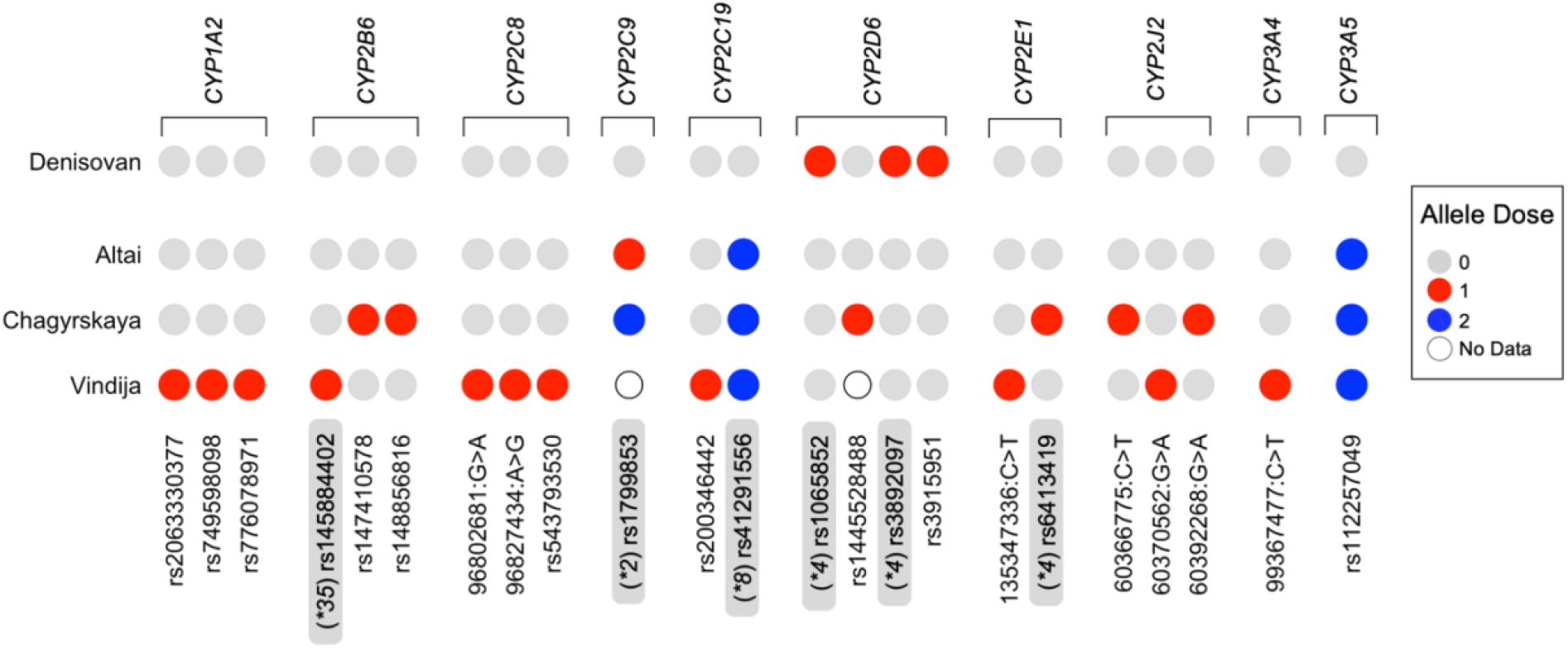
Function-altering variants in archaic individuals. Circle plot displaying the occurrence of potentially function-altering variants for each individual in the selected *CYP450* genes. Variants are identified with rsID, if no rsID number was identified, variants are considered potentially novel and the hg19 position is displayed along with mutation observed (position: reference allele > alternative allele). Variants that compose known star alleles are highlighted in gray and denoted with the respective star allele preceding the rsID. Allele dosage is categorized as the number of alternate alleles at that position. Un-filled circles indicate that no call was made at the given position for that individual. An allele dose of 1 indicates an SNV is heterozygous for the reference and alternate allele, and an allele dose of 2 indicates an SNV is homozygous for the alternate allele. No variants were identified for *CYP2A6*.

Of the potentially function-altering variants identified, 6 SNVs were diagnostic for star alleles (Figure 2 and 4, Table S5). *CYP2C9* rs1799853 is the core SNV for the *\*2* allele and was identified in the Altai and Chagyrskaya Neanderthal individuals, which is predicted to cause an intermediate metabolizer phenotype (or a slower metabolic rate) in modern humans. This SNV is also found globally in modern humans at less than 5% frequency, although European and admixed American populations from the 1000 Genomes Project Panel have higher frequencies of this allele, from 8-15% (Table S3). In *CYP2C19*, the primary SNV defining the *\*8* allele (rs41291556) was identified in the three Neanderthal individuals and confers a poor metabolizer status in modern humans (Figure 4, Table S5), although it is found at very low frequencies (0.5-1.5%) in a small number of modern human populations (Table S3). Finally, the two variants that compose the *CYP2D6*4* haplotype (rs3892097 and rs1065852) were identified as heterozygotes in the Denisovan individual and have been shown to confer an intermediate metabolizer status in modern humans (Figure 4). In addition to the two examples stated above, an exonic missense variant in *CYP2D6* was identified as heterozygous in a single Papuan (rs3915951) and was scored as likely to alter enzyme function (Figure 4, Table S3).

All 23 of the potentially function-altering alleles were considered derived according to the 1000 Genomes Project ^21^ ancestral allele and not identified in any of the chimpanzee samples ^27^ assessed (Table S5). Aside from the potentially function-altering variants identified, the majority of archaic variants present in non-African modern humans were either intronic (*n=*140) or exonic synonymous (*n=*4) mutations, making their impact on metabolism unclear (Figure 1C, Table S3).

### Introgression of archaic CYP450 alleles into modern humans

As shown in Table 1, each *CYP450* gene had a range of archaic variants found in modern human populations, from 2 shared SNVs (*CYP1A2*) to 64 shared SNVs (*CYP2C19*). This shared variation also varied extensively across global human populations, likely reflecting patterns of past interbreeding with diverse archaic populations and genetic drift ^9, 51^, but the shared archaic SNVs were largely at low frequency across populations (Figure 1C, Table S3). To confirm if the archaic SNVs identified in modern human populations for the *CYP450* genes corresponded to an archaic haplotype inherited from Neanderthals, we calculated sequence divergence between *CYP450* haplotypes using Haplostrips ^44^. Of the eleven *CYP450* genes studied here, the gene with the clearest evidence for haplotype sharing between Neanderthals and modern humans was *CYP2J2*, where a Neanderthal-like haplotype was present in all non-African populations at frequencies higher than 2%, with the highest frequencies found in admixed American populations (Figure 5, Figure S5). The Neanderthal-like *CYP2J2* haplotype in modern humans includes 8 SNVs that are found in Neanderthals, although the three SNVs that were predicted to be function-altering in the Neanderthal genome are absent from the modern human Neanderthal-like haplotypes (Figure 5, Table S3). Haplostrip plots for the ten additional *CYP450* genes are displayed in Figures S6 and S7.

**Figure 5.**
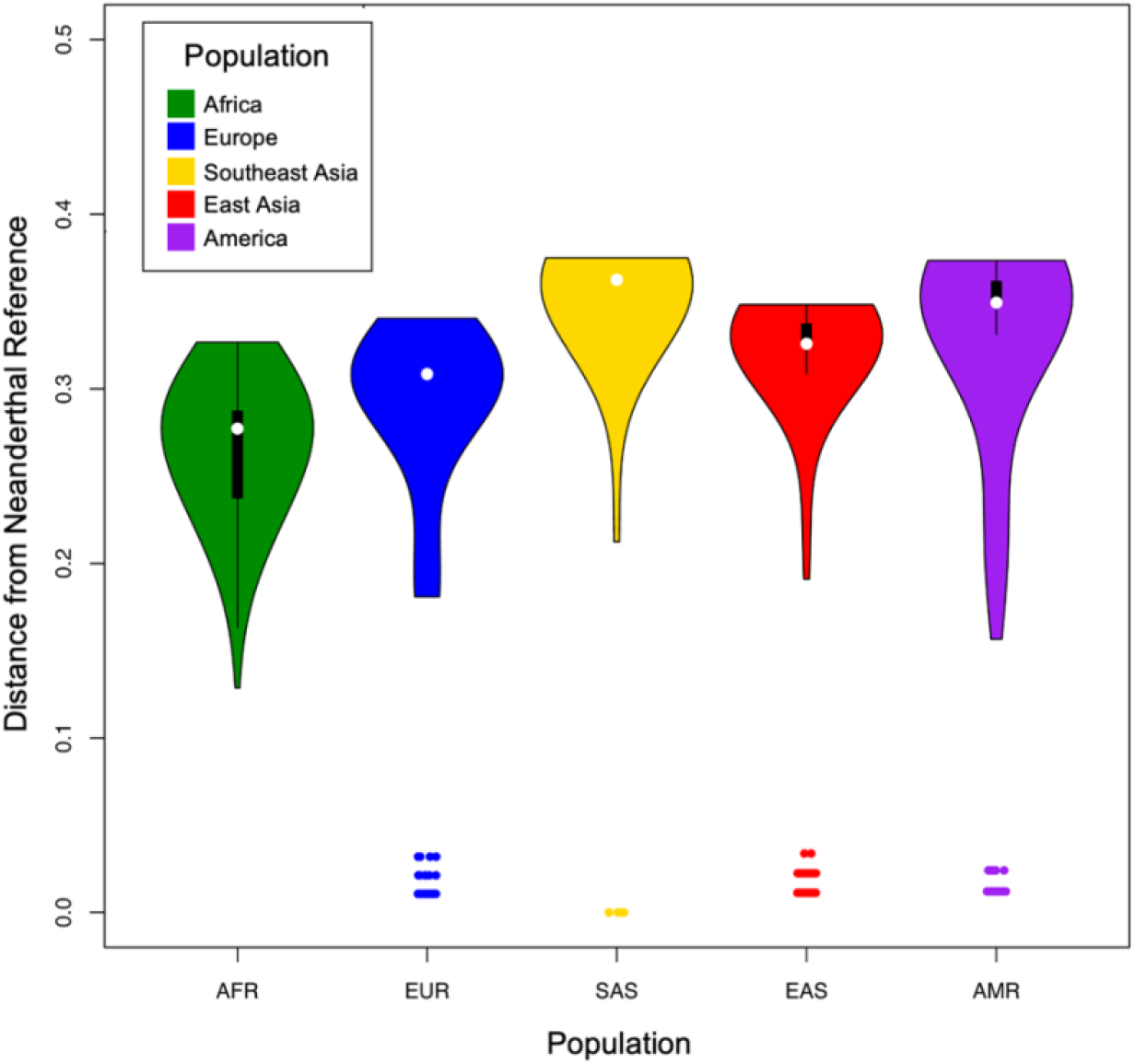
Haplostrips violin plot of *CYP2J2.* Haplotype distances at *CYP2J2* (hg19 1:60355979-60395470) in modern human populations relative to a Neanderthal reference haplotype (Vindija). The kernel density on each violin plots represents the abundance of individual haplotypes ordered by distance to the reference. Most human-like haplotypes sit at the same distance (furthest from the archaic reference), giving the violin plots a “flat top”. Outlier data points near lower distances represent modern human haplotypes carrying the Neanderthal-like *CYP2J2* in all populations except African populations.

**Table 1.**
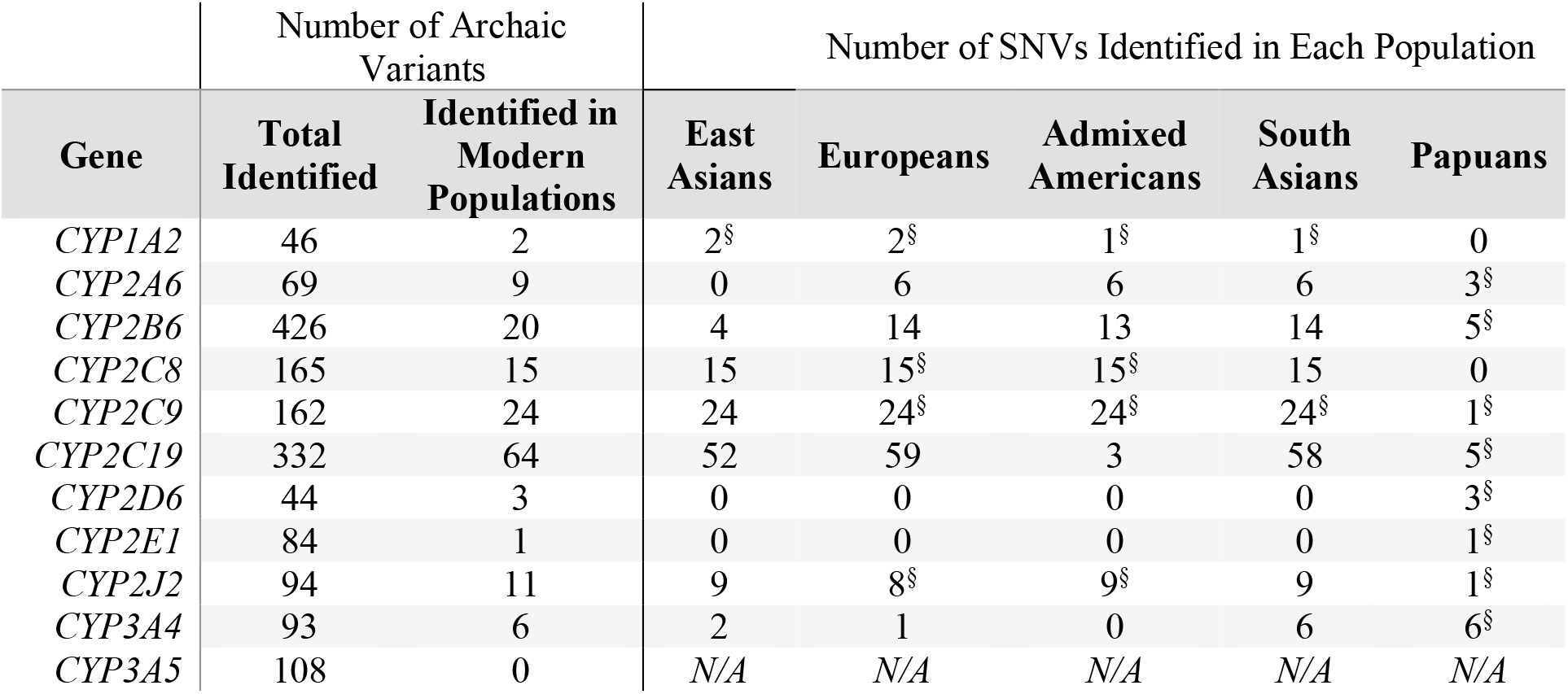
A summary of archaic variants identified in modern human populations for CYP450 genes. The number of SNVs found in each population is listed, along with the median allele frequencies for all SNVs identified in the populations in the region. “East Asians”, “Europeans”, “admixed Americans”, and “South Asians” refer to the populations from the 1000 Genomes dataset, while “Papuans” refers to the Simons Genome Diversity Project Papuan population. Variants of interest have been highlighted with a section sign (§).

## Discussion

We investigated a panel of *CYP450* genes in archaic individuals which are responsible for 75% of all drug metabolizing reactions and represent the bulk of drug metabolizing enzymes (DME) for which therapeutic recommendations exist ^17^. DMEs may be subject to evolutionary processes (e.g., mutation, drift, natural selection) because of their role as detoxifiers of xenobiotic substances, yet the insights from genetic variation in archaic genomes have not been characterized to date.

The majority of the *CYP450* genes in our panel conferred a normal/intermediate or unknown metabolizer phenotype (*CYP1A2*, *CYP2C8*, *CYP2C9*, *CYP2E1*, *CYP2J2*, *CYP3A4* and *CYP3A5*), consistent with previous studies showing strong purifying selection in pharmacogenes ^52^. Genes that were predicted to have a poor/slow (*CYP2A6*, *CYP2C19*) or ultrarapid (*CYP2B6*, *CYP2D6*) metabolizer status have also been known to have extreme variation in human populations ^53^ and may indicate instances of natural population variation or adaptation to the environment. However, it is important to note that the predicted phenotypes are based on modern human phenotypes, and it is unclear how the archaic-specific variation (particularly from the novel function-altering variants) would affect metabolism in archaic individuals. These variants of uncertain significance would benefit from further functional tests, individually as well as in the haplotypes found in archaic individuals, to measure both enzyme activity and protein abundance.

The distribution of archaic variants in the eleven *CYP450* genes we studied in modern populations is largely consistent with our general understanding of archaic introgression in human populations. For example, the majority of shared *CYP450* variants are Neanderthal in origin, which is consistent with the proportion of Neanderthal ancestry being significantly higher than the proportion of Denisovan ancestry for all populations except Melanesians ^9^. Papuans and other Melanesian populations have much more Denisovan ancestry than other populations ^26^ and may have had gene flow from a different Denisovan population than the one encountered by the ancestors of Eurasians ^54^, and thus, it is not surprising that they have a unique set of archaic alleles for the *CYP450* genes, many of which are Denisovan in origin.

Some of the variation shared between archaic and modern humans likely was present in the ancestral population to humans, Neanderthals, and Denisovans. Modern and archaic humans split relatively recently in evolutionary time and given the evolutionary constraints around pharmacogenetic enzyme structure and function, it is to be expected that Neanderthals and Denisovans would share similar diplotypes for humans for some of these genes. However, given the accumulation of neutral mutations over the half-million years of divergence between these species, particularly in introns and third codon positions, we are still able to distinguish between Neanderthal, Denisovan, and modern human haplotypes from this mutational background, even if functionally they present identical diplotypes.

### Evidence for archaic introgression in pharmacogenes

Given that pharmacogenes can be useful for adapting to novel environments ^2, 3^, we searched for evidence to suggest that some of the pharmacogene variation shared between archaic and modern humans were introduced through gene flow. Our primary criterion for identifying putatively archaic alleles was requiring that the archaic alleles were found in Africans at a frequency of less than 1% and found in at least one non-African population with a frequency higher than 1%. This method likely has an increased false-discovery rate (i.e., alleles shared between archaic and modern humans that are rare in Africa but are not introgressed) but provides a starting point for identifying potentially introgressed sites. While the goal of this study was to characterize variation rather than identify regions of selection or gene flow, here we suggest that some of the pharmacogenes may in fact show evidence for archaic introgression, which should be tested more rigorously for confirmation.

First, we infer archaic allele sharing between Europeans and Neanderthals in *CYP2C9.* The CYP2C9 enzyme is considered one of the most important DMEs in humans and responsible for metabolizing multiple classes of medications ^55, 56^ with substrates including non-steroidal anti-inflammatories ^57–60^, angiotensin II blockers (e.g. losartan) ^61^, S-warfarin ^57, 62^, phenytoin ^63^, and tolbutamide ^64^. In the *CYP2C9* gene, more than 60 haplotypes have been identified through the PharmVar consortium ^65, 66^. In this study, we identify multiple SNVs within *CYP2C9* that are shared between non-African populations and Neanderthals. Most interestingly, rs1799853, the causative SNV for the *CYP2C9*2* haplotype, is found in the Altai and Chagyrskaya Neanderthals and at frequencies of 8-15% in European populations, as well as in admixed American populations with a significant component of European ancestry ^67^. In modern humans, the *CYP2C9*2* variant has decreased enzyme activity compared to the reference haplotype ^68^, and negatively affects warfarin metabolism ^62^, increases the risk of hypoglycemia from sulfonylurea treatments ^69^ and increases the risk of overdose on phenytoin ^70^. However, it seems to also be protective against certain cancers ^71^. While some *CYP450* genetic variants (including *CYP2C19* and *CYP2D6* variants) have a global distribution consistent with the serial bottlenecks human populations experienced as they expanded out of Africa and into Eurasia, Oceania, and the Americas, the *CYP2C9* variants do not show the same correlation between genetic and geographic distance ^72^. If the *CYP2C9*2* variant was indeed introgressed from Neanderthals, the elevated frequency in European populations suggests that it may have been adaptive for processing certain xenobiotics that human populations were exposed to in Western Eurasia. However, at this time we cannot rule out that the elevated frequency of the Neanderthal allele in Europeans was the result of strong founder effect. Further exploration of this haplotype would be needed to determine whether this haplotype was introgressed and then targeted by positive selection.

Second, we identify a haplotype in *CYP2J2* that is found in all non-African populations but lacks the potentially function-altering variants found in Neanderthals, possibly suggesting that purifying selection has occurred for this gene. The CYP2J2 enzyme accounts for roughly 1-2% of hepatic CYP protein expression and is involved in the oxidation pathways of polyunsaturated fatty acids ^73^ and mediates the oxidation of drugs including ebastine ^74^, astemizole ^75, 76^, terfenadine ^76^, and ebastine ^77^, but also displays high expression in the lung, kidney, heart, placenta, salivary gland and skeletal muscle ^73, 78^. The majority of established *CYP2J2* polymorphisms occur at low frequencies, with the most common variant, *CYP2J2*7,* occurring at frequencies between 1-20% across global populations ^73, 79–84^. A comparison of haplotypes demonstrates that a Neanderthal-like haplotype is found in Eurasians and admixed American populations, with the highest frequency in the Americas (∼8%). Interestingly, the Neanderthal haplotype contains additional function-altering SNVs that are not found in modern humans. This suggests two possibilities: either the Neanderthal populations that admixed with modern humans did not carry these SNVs, or the function-altering SNVs were introduced to modern humans and then removed by strong purifying selection. Purifying selection has already removed much of the archaic ancestry in the modern human gene pool ^25, 26, 85–88^. This is likely because archaic humans had smaller population sizes, allowing for the accumulation of weakly deleterious alleles ^9, 51, 54, 89, 90^. After archaic and modern human admixture occurred, increased pressure from purifying selection would have removed archaic deleterious alleles, and it has been suggested that a significant portion of Neanderthal ancestry was removed from the modern human gene pool in just a few generations ^10, 91, 92^. The SNVs that remain in *CYP2J2* likely represent neutral variants that were maintained in the population or advantageous variants that may have been under positive selection. Further exploration of these archaic *CYP450* variants can help clarify the interplay between purifying selection removing deleterious mutations and positive selection retaining advantageous archaic *CYP450* alleles.

Finally, we identify a structural variant in *CYP2A6* that may have been introduced to modern humans through gene flow with Neanderthals. The *CYP2A6* gene is expressed mainly in the liver, and represents between 1% and 10% of the total liver CYP450 protein ^93^. The CYP2A6 enzyme metabolizes drugs and pro-carcinogenic compounds including tegafur ^94^, valproic acid ^95, 96^, and coumarin ^97^, and is the primary enzyme involved in nicotine metabolism ^98^. At present, more than 10 different allelic variants are known that cause absent or reduced enzyme activity ^99, 100^, suggesting that eliminating or reducing *CYP2A6* function is a recurring evolutionary strategy. The *CYP2A6* gene is located adjacent to the inactive *CYP2A7* gene and several allelic variants of *CYP2A6* have been created by unequal crossover and gene conversion reactions between these genes ^99^. The allele identified in Neanderthals is *CYP2A6*12*, a hybrid allele where exon 1 and 2 originated from *CYP2A7*, and exons 3–9 originated from *CYP2A6* (Figure 4). The *CYP2A6*12* allele causes a 50% reduction in CYP2A6 protein levels and a 40% decrease in CYP2A6 coumarin 7-hydroxylation activity ^99^. This hybrid *CYP2A6*12* allele has also been found at low frequencies in global populations, including African-American (0.4%), Canadian First Nation (0.5%), and Japanese (0.8%) individuals ^99^. In this study, we identify the *CYP2A6*12* hybrid allele in all three Neanderthal individuals as well as modern humans in Europe, Southeast Asia, and the Americas. This sharing of alleles, in particular the absence in unadmixed African populations, suggests that the human *CYP2A6*12* may have been inherited through human-Neanderthal introgression. There are alternate explanations that do not include archaic gene flow, such as incomplete lineage sorting (ILS) or parallel evolution. In the case of ILS, the unequal crossover event that created *CYP2A6*12* would have predated the split time of the modern human branch and the Neanderthal-Denisovan branch, ^101^, but this does not explain why the allele disappeared in Africans. Given the length of this haplotype (31.9 kb) and the regional recombination rate (0.77 cM/Mb ^45^), however, it is improbable that the 31.9 kb *CYP2A6*12* haplotype would have been maintained by ILS in both lineages (p=0.0014). ILS was additionally calculated for all eleven pharmacogenes investigated (Table S6). The final alternative to explain the prevalence of *CYP2A6*12* is that this hybrid allele evolved multiple times, independently in modern non-Africans and Neanderthals, in addition to the low frequency of *CYP2A6*12* introgressed from Neanderthals. It has been proposed that the slow metabolizing of some plant secondary metabolites may be adaptive as high levels of these toxins in human tissues may act as a deterrent to parasites ^102–104^, perhaps explaining why this hybrid allele may have evolved multiple times.

### Potential super-archaic introgression

For *CYP2B6*, we identified highly divergent haplotypes in the Vindija Neanderthal and a small number of African individuals (*n=*11). CYP2B6 is responsible for metabolizing drugs including the prodrug cyclophosphamide ^50, 105^; efavirenz, a non-nucleoside reverse transcriptase inhibitor ^50, 106, 107^; the antidepressant, bupropion ^50, 108, 109^; and ketamine ^110^. Genetic polymorphisms in *CYP2B6* have been identified to alter enzyme activity ^106, 111–114^ and there have been over 37 *CYP2B6* haplotypes identified ^66, 115^. In this study, we observe a haplotype in *CYP2B6* that is found in the Vindija Neanderthal and eleven modern African individuals and shows evidence of gene duplication. This duplicated haplotype is highly divergent from other archaic and modern human *CYP2B6* haplotypes, but the duplicated Neanderthal and human haplotypes are also distinct from one another (Supplemental Figure 4). The cause of the elevated SNV count in this duplicated haplotype is unclear. It is possible that the gene duplicate is non-functional, and the variation accumulated because selection has been relaxed, although there is no obvious evidence (i.e., a stop codon) to suggest that the enzyme’s function has been impaired. This gene duplication might also be an ancestral variant that has survived at low frequencies in modern human and Neanderthal populations, but this is unlikely given our ILS calculations for *CYP2B6* (p=0.02, see Supplemental Table 6). Finally, it is also possible that the duplicated haplotype is introgressed from an archaic hominin that is more divergent than Neanderthals and Denisovans, also known as a “super-archaic” human. There is evidence to suggest that Denisovans^21^ and Africans^6^ have small percentages of their genomes that were inherited from super-archaic humans, and *CYP2B6* was previously identified as a candidate region for super-archaic introgression in Africans^6^. Gene flow has also been inferred between Neanderthals and the ancestors of modern humans before either expanded outside of Africa ^116–118^. This implies that this gene duplication may have super-archaic origins. Future work on this haplotype’s function and evolutionary history may clarify its origin, and how it came to exist in both Neanderthals and modern humans.

## Limitations

Our findings suggest that Neanderthals and Denisovans have pharmacogene diplotypes that are similar to that of modern humans, and in some cases may reflect gene flow between archaic and modern humans. However, some limitations to the data must be noted. First, the archaic genomes are ancient DNA, which is degraded compared to modern DNA. This degradation often manifests as lower read depth, skewed allele balance, shorter read lengths, as well as transition point mutations (cytosine (C) → thymine (T)), which impact both mapping and sequencing accuracy. In order to distinguish true polymorphism at heterozygous sites from sequencing errors and DNA damage, we excluded base pairs that are represented in only a single read for each position in the BAM read alignment and removed any heterozygous calls that had an allelic read ratio less than 0.2 or greater than 0.8. The nature of ancient genomes resulted in low read depth for some star alleles that were called, but given this limitation, we also manually checked the sequencing reads for all diplotypes reported.

Additionally, haplotype calling on the archaic genomes required phasing of the archaic *CYP450* sequences using a human reference panel, and diplotype calls were made using current known human star alleles. While this is the best reference panel available, it is possible that phasing an archaic genome using modern human reference genomes will result in an inaccurate haplotype. However, the majority of SNVs were homozygous, which is consistent with the much lower heterozygosity of Neanderthals and Denisovans along the entire genome ^23–26^, and many of the heterozygous star allele calls were composed of only a single nucleotide variant, effectively deeming the phasing non-impactful. The concern that the archaic diplotypes may represent novel star alleles that are not shared with humans remains, and further functional validation of the novel potentially function-altering SNVs will be required to confirm the archaic diplotype calls.

We also used a novel read filtering method, resulting in a larger number of heterozygous sites compared to the variant call files that are usually used to represent the archaic genomes ^23–26^. The method used to make the previously generated variant call files likely used a more stringent method of filtering, resulting in more sites being called as homozygous. To ensure that our results were not biased by our variant calling method, we repeated all analyses (including calling star alleles, identifying archaic SNVs in modern humans, and comparing human and archaic haplotypes) using the original variant call files, and found that while the number of identified SNVs is smaller, the patterns identified (such as the divergent *CYP2B6* haplotype in the Vindija and allele sharing between archaic and modern humans) remain.

## Conclusions

Understanding the impact of archaic variation on modern human health is still in its early stages. While a few genes have been identified for which the archaic alleles play an important role in human adaptation to their environment, such as *EPAS1*, the vast contribution of archaic ancestry to human fitness is less clear. For example, modern humans carry archaic variants of medically important genes such as ABO blood group antigens, but their impact to human health remains unknown ^119^. Our results suggest that interactions between modern and archaic humans may have resulted in the introduction of novel *CYP450* variants to modern human populations, helping them adapt to novel environments as they expanded out of Africa. Important insights will continue to emerge from careful inspection of pharmacologically relevant and highly studied genes such as the *CYP450* genes investigated here. We look forward to identifying archaic *CYP450* alleles inherited by modern humans that have been adaptive in historically underrepresented populations in genomic research such as Indigenous American populations, which have uniquely adapted to numerous novel environments and have undergone multiple genetic bottlenecks due to colonization and disease.

## Data and Code availability

The code generated during this study is available on GitHub (https://github.com/the-claw-lab/aDNA_PGx_2021 and https://github.com/kelsey-witt/archaic_pgx).

## Author contributions

THW, KEW, SBL, RSM, EHS, FV, and KGC designed research, project conception and development. THW, KEW, SBL, RSM, EHS, FV, and KGC conducted research, formal analysis, and manuscript writing – original draft. THW, KEW, SBL, RSM, EHS, FV, and KGC performed manuscript review and editing. THW, KEW, SBL, RSM, EHS, FV, and KGC provided resources and study oversight. THW, KEW, FV, and KGC have primary responsibility for final content. All authors read and approved final manuscript

## Acknowledgements

We would like to thank George Perry and Omer Gokcumen for comments on early versions of our manuscript, and anonymous reviewers for their feedback.

