## Supplemental Figures for "Pharmacogenetic variation in Neanderthals and Denisovans and implications for human health and response to medications"

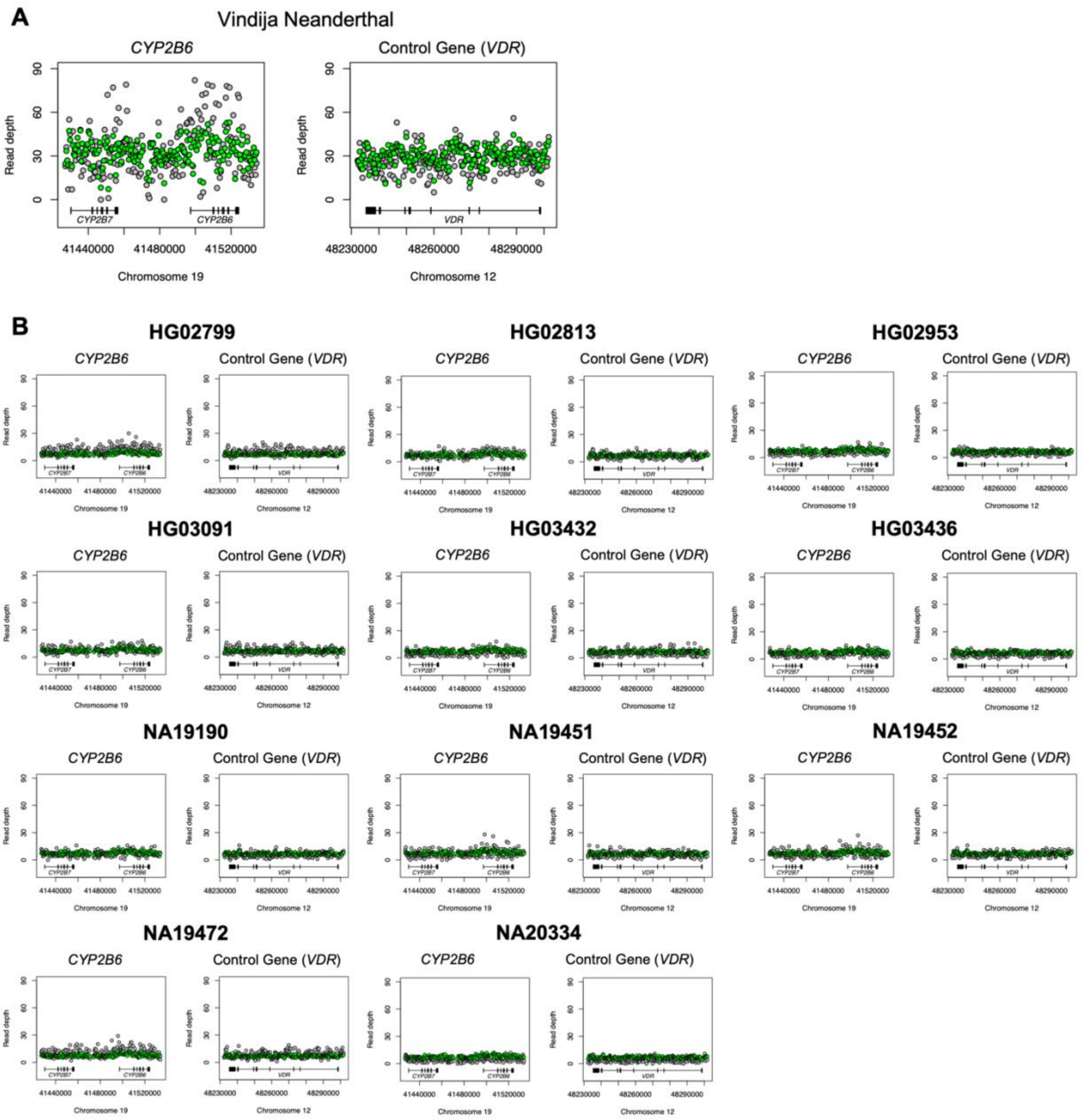


**Figure S1.** ***CYP2B6*** **read depth plots.** Read depth plots for the (A) Vindija Neanderthal and (B) 11 divergent African modern humans. Plots displayed as the read depth calculated from the BAM files for *CYP2B6* / *CYP2B7* (pseudo-gene, left plot) and the *VDR* control gene region (right plot). Hg19 genetic coordinates and chromosome are presented on the *x axis*.

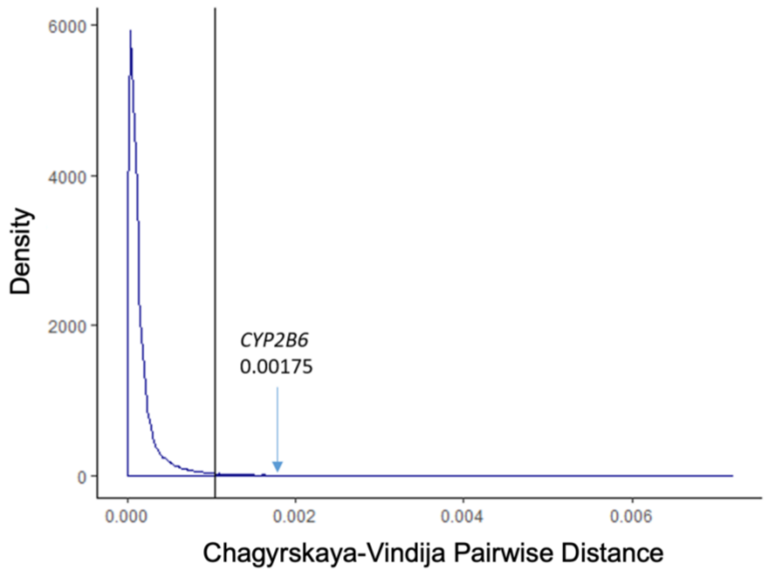


**Figure S2. Pairwise distance distribution between the Vindija and Chagyrskaya Neanderthals.** The distribution of pairwise distances of the Chagyrskaya and Vindija genomes for 29.1-kb windows (*CYP2B6* gene length) across the genome. The cutoff for the top 1% of values is labeled with a vertical line, and the pairwise distance for the *CYP2B6* gene is labelled and denoted with an arrow.

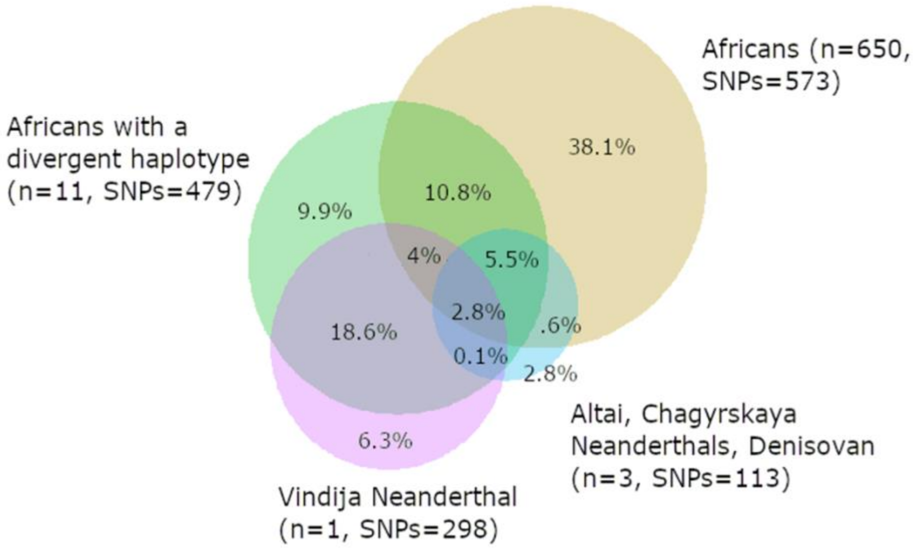


**Figure S3. Shared SNVs between modern and archaic humans in *CYP2B6*.** A Venn diagram that illustrates the overlap in alternate alleles found in *CYP2B6* for modern and archaic humans. Here, the eleven African individuals with a divergent haplotype are considered separately from the rest of the African individuals (simply labeled here as ‘Africans’), and the Vindija Neanderthal is considered separately from other archaic humans. For each population, the sample and total number of variable sites (SNVs) is noted. For each portion of the Venn diagram the percentage denotes the total percentage of variable sites in the *CYP2B6* gene that are shared with the included populations.

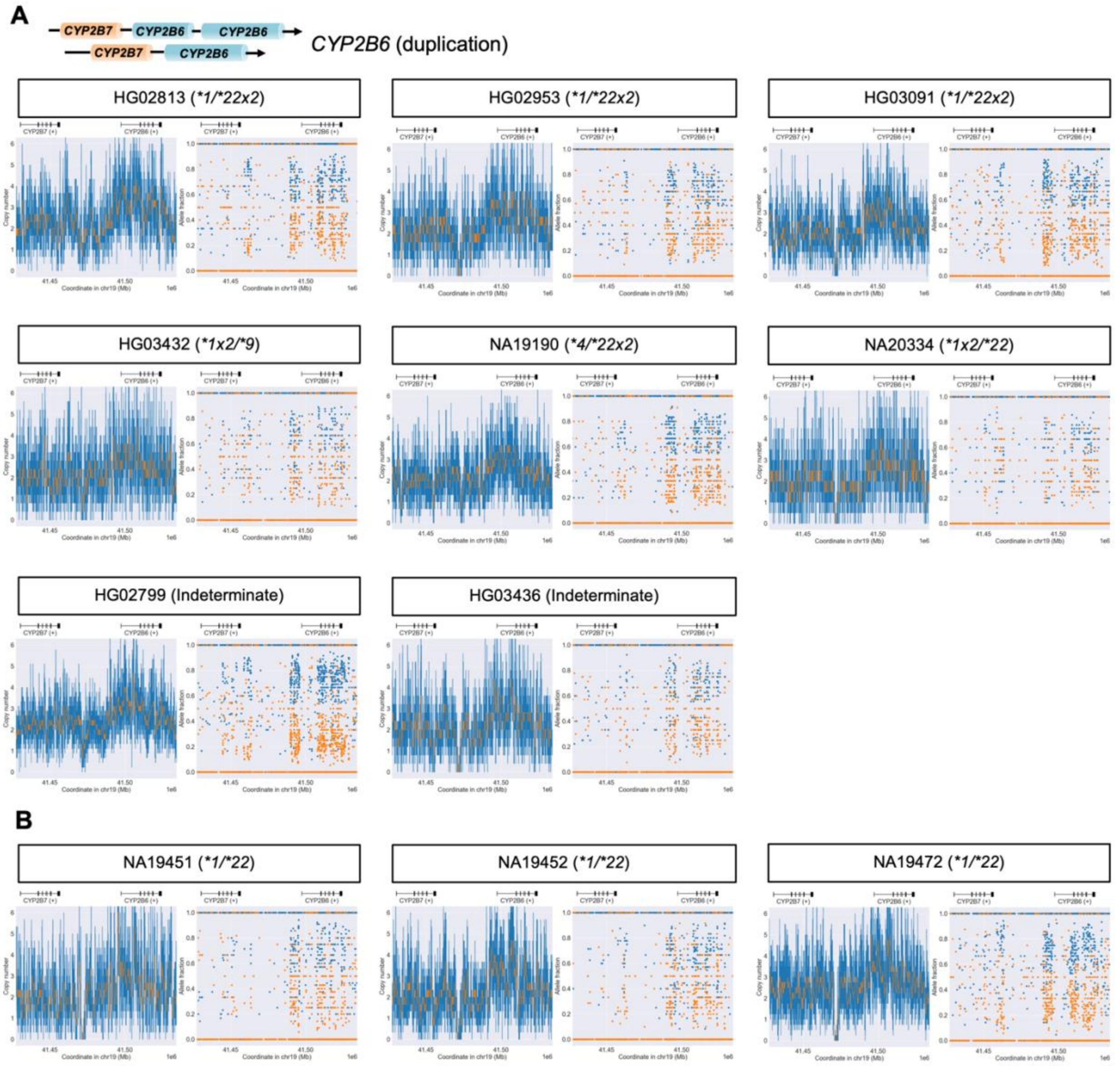


**Figure S4. Structural variant plots for *CYP2B6* in the 11 divergent African individuals.** Copy number variation plots (left) demonstrating the structural variation and allele fraction plots (right) showing allelic fraction for each divergent African individual. (A) *CYP2B6* gene duplication was identified in 8/11 individuals, and (B) manual inspection in the three other modern African individuals suggests elevated copy number as well. Copy number plots are estimated based on read data (displayed as the copy number normalized to the VDR control gene region) with the orange line indicating the copy number assessment for each gene. Allele fraction plots display the allele frequency of each variant for the two haplotypes, which demonstrates the allelic decomposition after identifying structural variation. A schematic for *CYP2B6* gene duplication is displayed above panel A with an arrow indicating the direction of transcription. The hg19 genetic coordinates are presented on the *x axis* with the gene regions indicated directly above each plot.

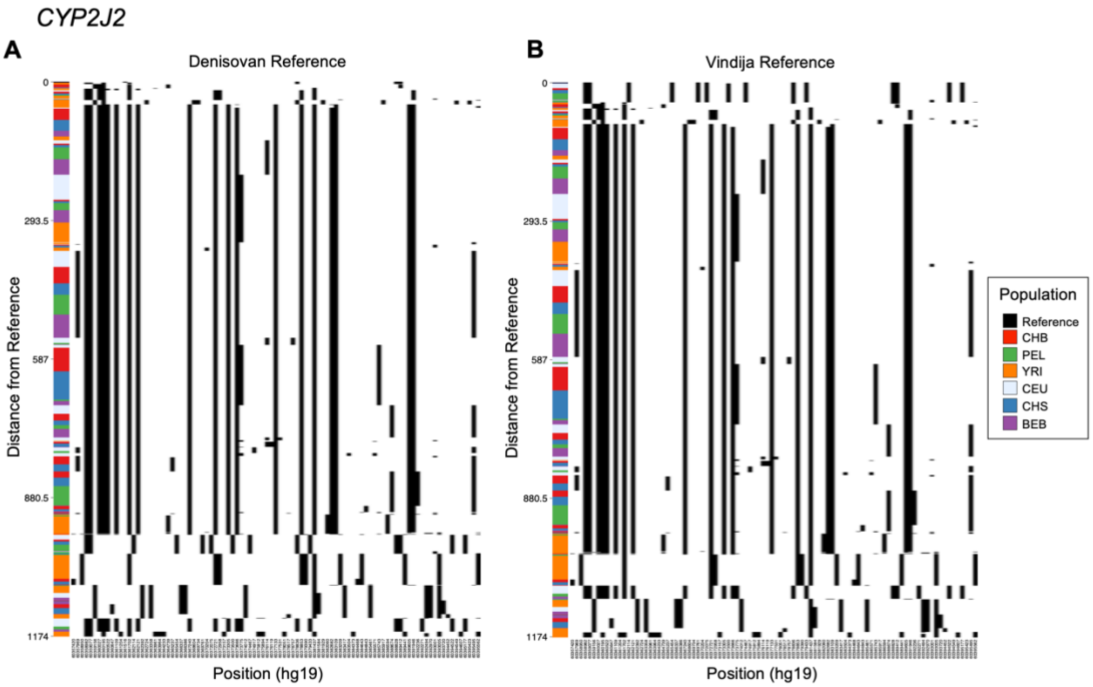


**Figure S5. Haplostrips plot for *CYP2J2*.** Visualization of the sequence divergence between gene haplotypes from individuals from the 1000 Genomes Project relative to the (A) Denisovan reference haplotype and (B) Vindija Neanderthal reference haplotype. Each haplotype is represented as a horizontal line, ordered by sequence divergence relative to the reference haplotype. Colors along the *y axis* represent the super population for each haplotype in the haplostrip.

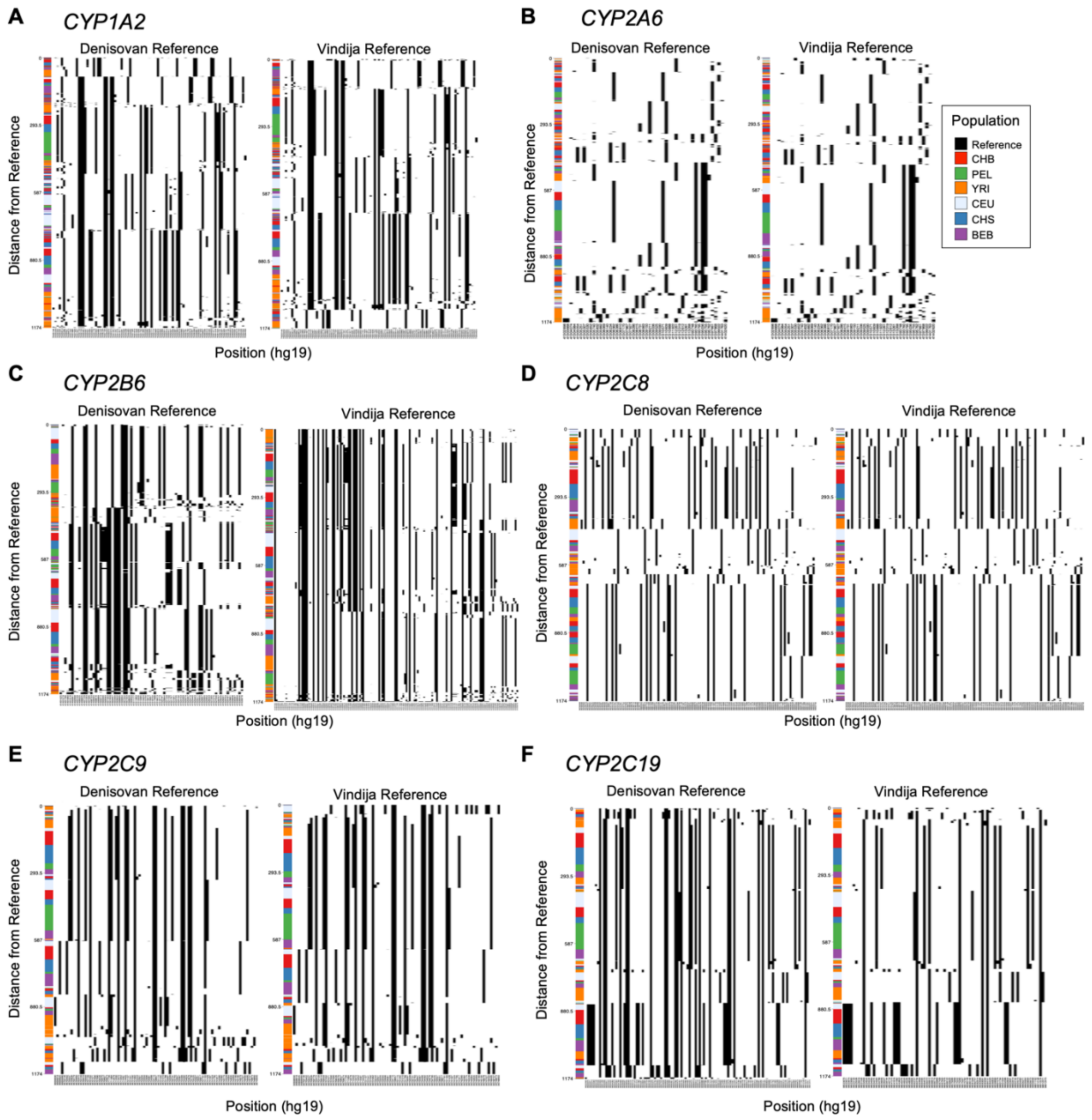


**Figure S6. Haplostrips plots for *CYP450* genes.** Visualization of the sequence divergence between gene haplotypes from individuals from the 1000 Genomes Project relative to the Denisovan reference haplotype (left) and Vindija Neanderthal reference haplotype (right) for (A) *CYP1A2,* (B) *CYP2A6,* (C) *CYP2B6,* (D) *CYP2C8,* (E) *CYP2C9,* and (F) *CYP2C19*. Each haplotype is represented as a horizontal line, ordered by sequence divergence relative to the reference haplotype. Colors along the *y axis* represent the super population for each haplotype in the haplostrip.

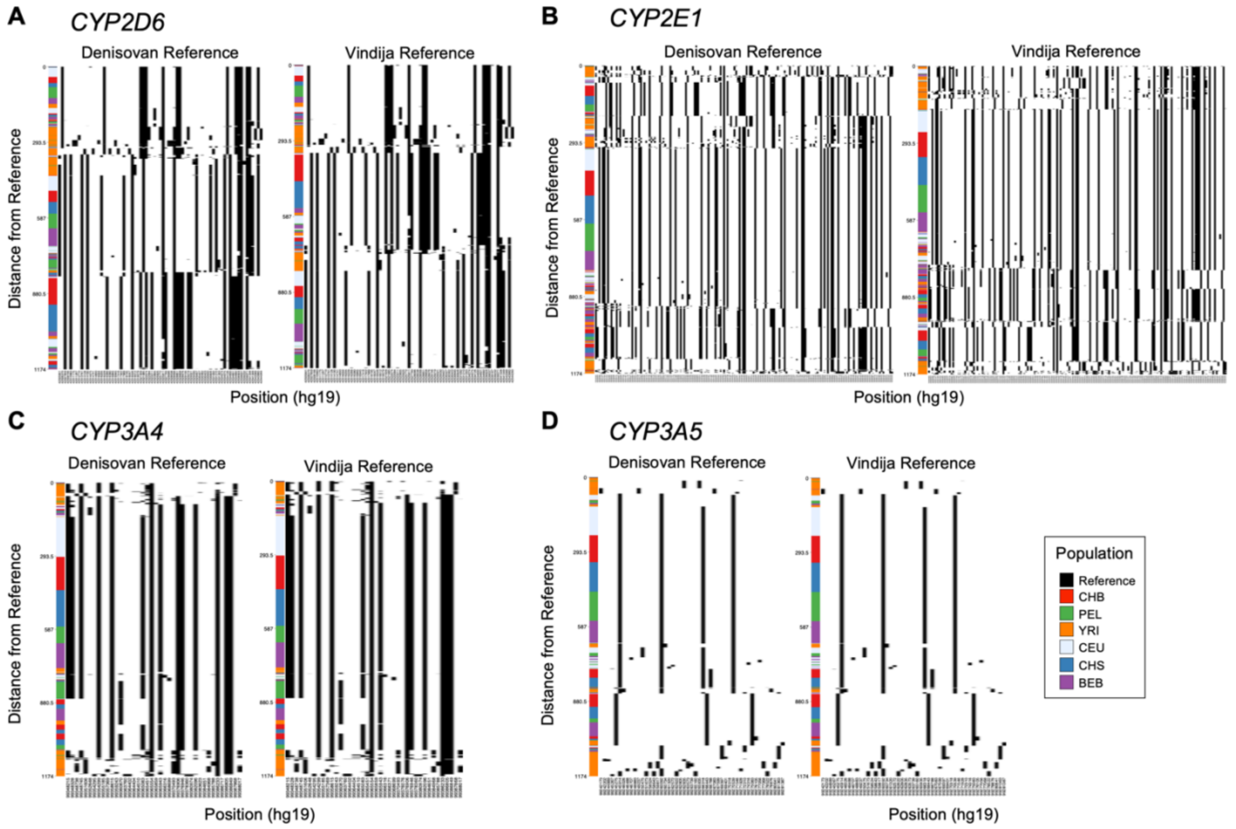


**Figure S7. Haplostrips plots for *CYP450* genes.** Visualization of the sequence divergence between gene haplotypes from individuals from the 1000 Genomes Project relative to the Denisovan reference haplotype (left) and Vindija Neanderthal reference haplotype (right) for (A) *CYP2D6,* (B) *CYP2E1,* (C) *CYP3A4,* and (D) *CYP3A5*. Each haplotype is represented as a horizontal line, ordered by sequence divergence relative to the reference haplotype. Colors along the *y axis* represent the super population for each haplotype in the haplostrip.
